# The Pax transcription factor EGL-38 links EGFR signaling to assembly of a cell-type specific apical extracellular matrix in the *Caenorhabditis elegans* vulva

**DOI:** 10.1101/2024.09.04.611291

**Authors:** Helen F. Schmidt, Chelsea B. Darwin, Meera V. Sundaram

## Abstract

The surface of epithelial tissues is covered by an apical extracellular matrix (aECM). The aECMs of different tissues have distinct compositions to serve distinct functions, yet how a particular cell type assembles the proper aECM is not well understood. We used the cell-type specific matrix of the *C. elegans* vulva to investigate the connection between cell identity and matrix assembly. The vulva is an epithelial tube composed of seven cell types descending from EGFR/Ras-dependent (1°) and Notch-dependent (2°) lineages. Vulva aECM contains multiple Zona Pellucida domain (ZP) proteins, which are a common component of aECMs across life. ZP proteins LET-653 and CUTL-18 assemble on 1° cell surfaces, while NOAH-1 assembles on a subset of 2° surfaces. All three ZP genes are broadly transcribed, indicating that cell-type specific ZP assembly must be determined by features of the destination cell surface. The paired box (Pax) transcription factor EGL-38 promotes assembly of 1° matrix and prevents inappropriate assembly of 2° matrix, suggesting that EGL-38 promotes expression of one or more ZP matrix organizers. Our results connect the known signaling pathways and various downstream effectors to EGL-38/Pax expression and the ZP matrix component of vulva cell fate execution. We propose that dedicated transcriptional networks may contribute to cell-appropriate assembly of aECM in many epithelial organs.

**Highlights:**

- *C. elegans* vulva apical extracellular matrix is cell-type specific
- Broadly transcribed Zona Pellucida domain proteins assemble in specific matrices
- The Pax2/5/8 homolog EGL-38 promotes assembly of the 1° vulva cell matrix
- EGL-38 expression and 1° cell matrix assembly depend on EGFR signaling

## Introduction

A common feature of epithelial cells is an apical extracellular matrix (aECM) which covers their exposed surfaces. Molecularly distinct aECMs such as the cornea tear film, the vascular glycocalyx, the gut mucin lining, and the cochlea tectorial membrane have distinct roles in the protection, function, and/or development of the respective tissue (Butler et al., 2020; Goodyear and Richardson, 2018; Gustafsson and Johansson, 2022; Martinez-Carrasco et al., 2021). Whether it is produced by the epithelial cells themselves or secreted by specialized cells such as goblet cells or glands (Gustafsson and Johansson, 2022; Martinez-Carrasco et al., 2021), assembling and maintaining the proper aECM is an important aspect of epithelial cell function. Defects in the aECM contribute to many human diseases, including allergic conjunctivitis, atherosclerosis, deafness, and irritable bowel disease (Butler et al., 2020; Goodyear and Richardson, 2018; Gustafsson and Johansson, 2022; Martinez-Carrasco et al., 2021). How particular epithelial cells control assembly of the correct matrix components on their apical surfaces is not well understood, particularly given that relevant matrix factors sometimes originate from other tissues or cell types.

*C. elegans* is an excellent model to study the connection of epithelial cell identity to its aECM. It has an invariant lineage, with at least 30 different epithelial cell types, each of which can be viewed at single cell resolution (Sulston and Horvitz, 1977). Additionally, the aECMs covering these epithelia contain many types of conserved proteins, including collagens and zona pellucida domain (ZP) proteins (Sundaram and Pujol, 2024). These matrix proteins are arranged in distinct domains on the surface of different epithelial cell types and are dynamic as the worm develops (Adams et al., 2023; Cohen et al., 2020b; Katz et al., 2022; Serra et al., 2024). The worm’s epidermis and interfacial tubes are covered by a collagenous matrix called the cuticle. At each larval stage, the worm builds a fresh cuticle and molts the old one. Construction of the cuticle is influenced by the pre-cuticle, a ZP-rich aECM that is endocytosed prior to molting (Birnbaum et al., 2023; Katz et al., 2022; Serra et al., 2024; Vuong-Brender et al., 2017). These dynamic, tissue- and cell type-specific aECMs present multiple avenues to study the connection between cell identity and matrix.

Here we focus on a particular epithelial tube, the vulva, and its aECM. The vulva connects the uterus to the external epithelia and is used for egg laying and mating. There are seven vulva cell types (vulA, B1, B2, C, D, E and F), with two or four cells of each type for a total of 22 cells (Figure 1A, B) (Sharma-Kishore et al., 1999; Sulston and Horvitz, 1977). During the final (L4) larval stage, these cells invaginate to form a lumen and cells of the same type meet and in some cases fuse with each other to produce seven stacked rings (Sharma-Kishore et al., 1999) (Figure 1A). The vulva aECM is dynamic during development. The vulva lumen initially fills with a chondroitin proteoglycan-rich matrix promoting inflation (Herman et al., 1999; Hwang et al., 2003). Multiple pre-cuticle factors then assemble on the surface of different cell types and contribute to shaping of the developing vulva (Cohen et al., 2020b). This pre-cuticle is ultimately replaced by the final adult cuticle.

**Figure 1:**
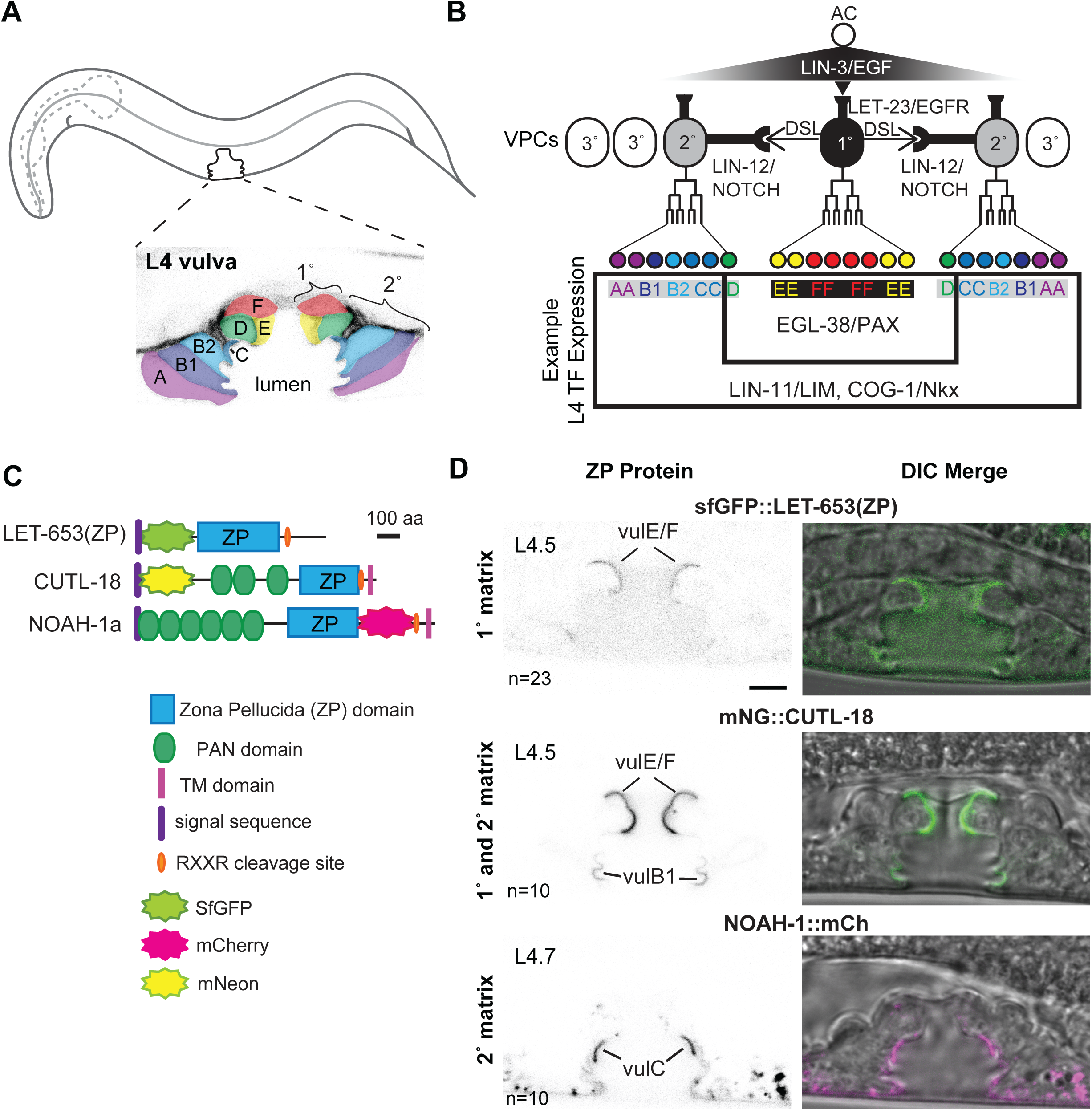
Zona Pellucida domain (ZP) protein matrix is cell type-specific A) Position of the vulva and L4.5 stage vulva cells visualized with the membrane marker MIG-2::GFP (*muIs28*) (Honigberg and Kenyon, 2000). The seven vulva cell types (vulA-F) are highlighted by color. B) Specification of vulva cells. The anchor cell (AC) releases a gradient of LIN-3/EGF to the six equipotent vulva precursor cells (VPCs-ovals). The closest cell receives the strongest EGF signal and adopts the primary (1°) vulva fate, dividing to give rise to vulE/F. The 1° VPC also expresses Delta/Serrate/LAG-2 (DSL) ligands to activate LIN-12/Notch signaling in neighboring VPCs and specify the secondary (2°) vulva fate, which gives rise to vulA/B1/B2/C/D. Transcription factors (TFs) controlling aspects of each cell fate are sometimes broadly expressed in the vulva (e.g. LIN-11/LIM, COG-1/Nkx), or expressed in specific vulva cells (e.g. EGL-38/Pax). C) ZP protein schematics. The LET-653(ZP) fusion consists of the isolated ZP domain and was expressed from a transgene (*csIs66*) (Cohen et al., 2019); full-length LET-653 has multiple PAN domains at its C-terminus. NOAH-1 and CUTL-18 fusions were each generated by CRISPR-Cas9 genome editing and expressed from the endogenous locus. D) ZP proteins localize to the matrix of specific vulva cells. LET-653(ZP) and CUTL-18 are 1° matrix components on vulE/F. CUTL-18 is also a 2° matrix component on vulB1/B2. NOAH-1 is a 2°-specific matrix component, enriched on vulC. Images are medial confocal slices at the indicated L4 stages. Scale bar 5 µm.

The vulva has been extensively studied as a model for signaling and cell type specification (Shin and Reiner, 2018). Vulva cells arise from a group of six equipotent vulva precursor cells (VPCs) that adopt different fates in response to signaling mediated by the Epidermal Growth Factor receptor (EGFR)-Ras-ERK and Notch pathways (Figure 1B). The anchor cell releases a LIN-3/EGF signal received by LET-23/EGFR on the nearest VPC, causing it to adopt the primary (1°) cell fate. The 1° VPC then produces Delta/Serrate/Lag2 (DSL) ligands and signals through LIN-12/Notch for the two adjacent VPCs to adopt the secondary (2°) cell fate (Figure 1B). These three VPCs then divide in a stereotyped pattern to produce the 1° lineage-derived cells vulE and F and the 2° lineage derived cells vulA, B1, B2, C, and D (Figure 1B). The additional VPCs are not induced to become vulval and instead divide once and fuse with the hypodermis (3° fate). Mutations in signaling genes cause VPC cell fate transformations and either loss or gain of 1° and/or 2° vulva lineages, resulting in Vulvaless (Vul) or Multivulva (Muv) phenotypes.

While the specification of VPC lineage fate is well understood, it is less clear how each fate is executed and how the expression of downstream genes is controlled in each of the 7 distinct cell types. Many transcription factors are expressed in the vulva (Figure 1B), but unlike signaling mutants, these transcription factor mutants don’t have clean cell fate transformations but instead lose (or mis-express) only a subset of known cell fate markers in one or more cell types (Fernandes and Sternberg, 2007; Gupta et al., 2012; Inoue et al., 2005). This suggests that each vulva cell type’s identity is controlled by a combination of transcription factors that work together.

EGL-38, a Pax2/5/8 related transcription factor, controls some aspects of vulE/F development (Chamberlin et al., 1997). EGL-38 is expressed in the 1° lineage from soon after VPC specification through mid-L4, as well as faintly in vulD (Mok et al., 2015; Webb Chasser et al., 2019). *egl-38* mutants have a normal vulva cell lineage but defects in vulE/F lumen shape, consistent with EGL-38 playing a role in some aspect of fate execution (Chamberlin et al., 1997; Trent et al., 1983). One target of EGL-38 in the vulva is *lin-3*/EGF, which is required for signaling to specify uterine uv1 cells (Chang et al., 1999; Rajakumar and Chamberlin, 2007). EGL-38 is not required for proper vulE or vulF cell fusion or for expression of other known vulE/F markers such as *zmp-1* and *bam-2* (Inoue et al., 2005; Rajakumar and Chamberlin, 2007), but it does function redundantly with other transcription factors to prevent vulE/F from expressing inappropriate 2° fate markers (Fernandes and Sternberg, 2007). Further study has been limited by the few known markers of vulva cell type biology.

Here we use aECM as a marker to assess the contribution of multiple transcription factors to vulva cell identity. We find that ZP protein gene transcription patterns are much broader than the observed cell-type specific patterns of ZP protein localization, indicating that each cell recruits the proper matrix to its surface. Using mutants in EGFR effector genes, we find that vulva cells’ matrix identity is separable from their identity as assessed by lineage. We find EGL-38 is required for the aECM identity of vulE/F. Overall these data suggest a model where EGL-38 promotes the transcription of one or more matrix organizers that promote the assembly of the proper ZP matrix.

## Results

### Pre-cuticle and cuticle ZP proteins show cell-type specific localization patterns in the vulva

First identified as components of the mammalian egg coat, ZP proteins are common components of aECMs across organisms (Plaza et al., 2010). ZP domains are polymerization modules whose assembly can be influenced by proteolysis and/or interactions with matrix partners (Figure 1C) (Bokhove and Jovine, 2018; Jovine et al., 2002)). In *C. elegans*, ZP proteins are required to shape the body and multiple interfacial tubes, including the vulva (Gill et al., 2016; Sapio et al., 2005; Vuong-Brender et al., 2017). We previously found two pre-cuticle ZP proteins (LET-653 and NOAH-1) were closely associated with the aECM on specific vulva cell surfaces (Cohen et al., 2020b; Gill et al., 2016), making them ideal candidates to study the connection between cell identity and aECM. In addition, we used published single cell RNA sequencing data (Taylor et al., 2021) to identify CUTL-18 as an additional ZP protein likely present in the vulva and other L4 interfacial tubes. Like LET-653 and NOAH-1, CUTL-18 has multiple PAN domains followed by a ZP domain. Like NOAH-1, but unlike LET-653, CUTL-18 has a C-terminal transmembrane domain (Figure 1C). All three proteins have predicted cleavage sites immediately following their ZP domains (Figure 1C).

We observed the localization of these three vulva-expressed ZP domain proteins using fluorescently-tagged proteins expressed from transgenes (LET-653) or from the endogenous loci (NOAH-1 and CUTL-18) (Figure 1C). Consistent with previous findings, LET-653(ZP) is enriched on the surface of the 1° lineage descendants vulE/F at mid-L4 (Gill et al., 2016, Figure 1D). NOAH-1 is present on the surface of all 2° lineage descendants, but strongly enriched on a matrix “spike” on vulC in later (L4.7/8) L4 larvae (Cohen et al., 2020b) (Figure 1D).

We found CUTL-18 is an additional component of the 1°matrix and is also present faintly on the 2°-derived cells vulB1/B2 (Figure 1D). Unlike the pre-cuticle components LET-653 and NOAH-1 (Cohen et al., 2020b; Vuong-Brender et al., 2017), CUTL-18 persists as a component of the adult cuticle where it lines the adult vulva, excretory duct, and the junction between the rectum and intestine (Supplementary Figure 1). CUTL-18 was not observed in the body cuticle and therefore is a tube-specific cuticle component. We conclude that vulva cuticle assembly begins as early as the L4.4 stage, concurrent with pre-cuticle assembly, and that both pre-cuticle and cuticle ZP proteins have cell-type specific assembly patterns.

### LET-653 and CUTL-18 independently localize to 1° matrix

ZP domains can interact with each other and, in some cases, ZP proteins depend on each other for proper secretion or assembly (Bokhove et al., 2016; Chu and Hayashi, 2021; Drees et al., 2023; Ghosh and Treisman, 2024; Vuong-Brender et al., 2017). Since CUTL-18 and LET-653 both localize to the 1° cell matrix, we tested whether sfGFP::LET-653(ZP) or mNG::CUTL-18 localize normally in the vulva of worms mutant for the other gene. For *let-653*, we used a null mutation (*cs178*) rescued to viability with an excretory tube-specific transgene (Gill et al., 2016). For *cutl-18*, we used a splice acceptor mutation (*gk516154*) predicted to lead to a frameshift and premature stop (Thompson et al., 2013). Neither ZP protein gene was required for the other protein’s localization (Supplementary Figure 2).

### Specificity of ZP protein localization is not determined by patterns of ZP gene transcription

We tested whether the cell-type specific localization of these ZP proteins could be explained by cell-type specific transcription. To capture the full transcriptional regulatory context of the ZP protein genes, we generated endogenous reporters by CRISPR/Cas9 genome editing. At each locus, mCherry::HIS-44(H2B) was expressed in an operon with the ZP protein gene using the SL2 spliced leader sequence (Figure 2A). Given the irregular three-dimensional shape of vulva cells, these nuclear-localized reporters allowed clearer cell type identification than cytoplasmic reporters.

**Figure 2:**
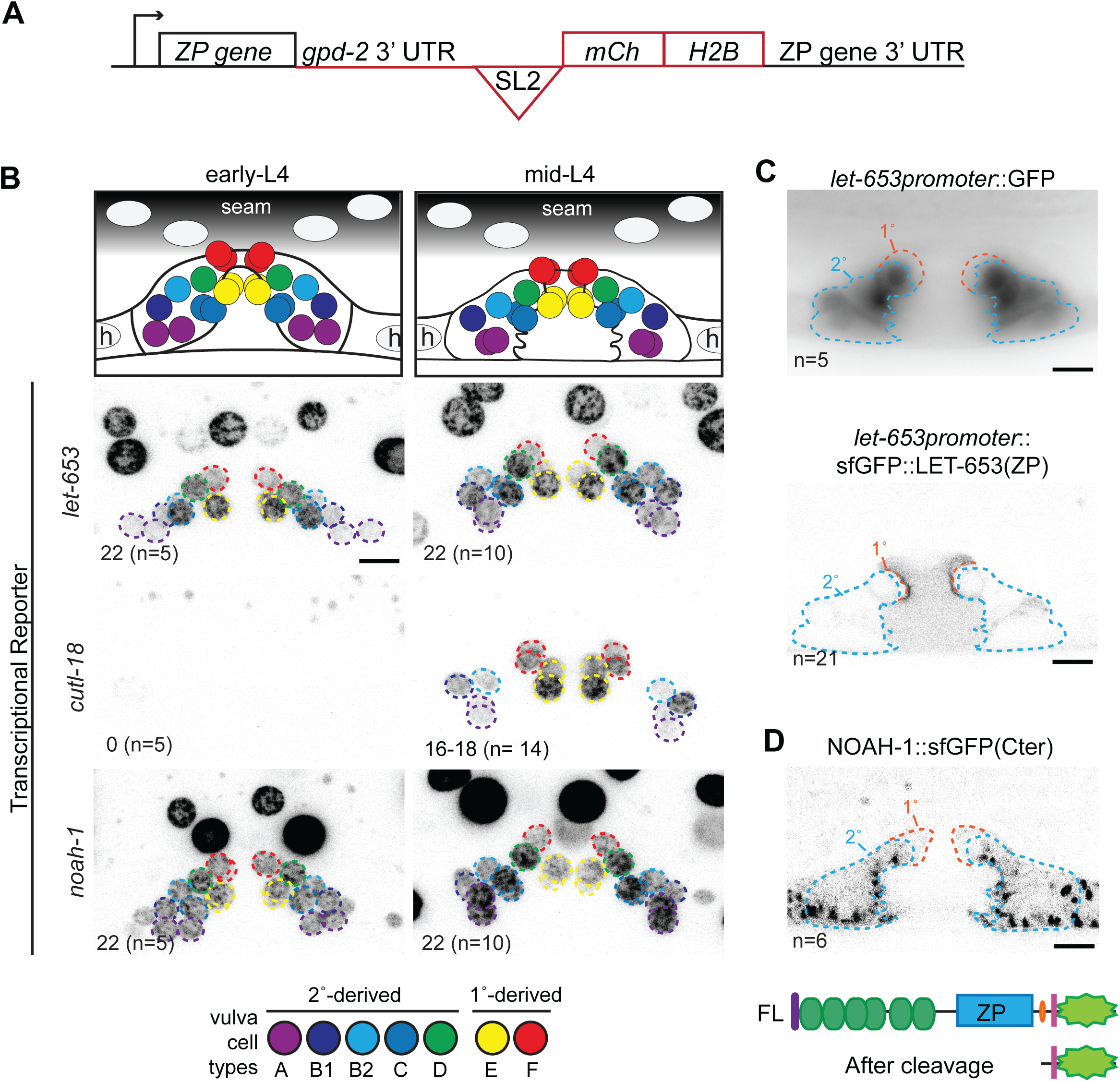
ZP protein localization is not explained by their transcription patterns A. Schematic of transcriptional reporters expressed from the endogenous loci of ZP protein genes. Sequence inserted immediately after stop codon of each ZP protein gene is outlined in red. The *gpd-2* 3’UTR and trans-spliced leader sequence (SL2) allow the ZP protein and mCh::HIS-44 (H2B) to be expressed as an operon. B. ZP protein gene expression in early-(L4.1) and mid-(L4.4/L4.5) L4 larvae. Top: diagram of approximate position of each vulva nucleus at the indicated stages. Below: Maximum projections of confocal Z stacks through entire vulva of worms expressing the indicated transcriptional reporter. Vulva nuclei are outlined according to the color code below. Lower left: Number of mCherry positive nuclei, (n= number of worms). The *cutl-18* transcriptional reporter varied slightly at the mid-L4 stage; it was present in vulA,B1,B2,E, and F (16 total nuclei) in all worms, and also faintly in vulD (18 total nuclei) in 11/14 worms. C. LET-653(ZP) assembles on the surface of 1° cells in absence of its transcription in 1° cells. Mid-L4 vulvas. Top: epifluorescent image of cytoplasmic GFP expressed from a 2233bp *let-653* promoter fragment (Hunt-Newbury et al., 2007); GFP is present in 2° cells and faintly in hyp7. Bottom: medial confocal slice of sfGFP::LET-653(ZP) expressed from the same promoter fragment is present inside 2° cells (orange outline) and assembles on 1° cell surfaces. D. The NOAH-1 C-terminus (Cter) is not secreted or cell-type specific. Top: Since NOAH-1::sfGFP(Cter) is predicted to be cleaved N-terminal to the transmembrane domain, the sfGFP remains inside the cells that produced it (Vuong-Brender et al., 2017). Medial confocal slice of L4.5 stage vulva. NOAH-1 C-terminus forms large puncta in all 2° vulva cells and hyp7. Consistent with the transcriptional reporter, NOAH-1::sfGFP(Cter) signal is not distinct between vulC and other 2° cells or hyp7. Below: Schematic of NOAH-1 with sfGFP at its C-terminus, full length (FL), and predicted product after cleavage. Domain symbols as are in Figure 1. All scale bars 5µm.

The endogenous *let-653* and *noah-1* reporters were transcribed broadly in external epithelia, including in all vulva cells by the beginning of the L4 stage (Figure 2B). In contrast, the *cutl-18* reporter appeared tube-specific; in the vulva it was first detected in mid L4 (L4.4/5) and was transcribed in most vulva cells but not vulC (Figure 2B). All three ZP protein genes were transcribed in multiple cell types that did not have that protein in their aECM. Consistent with broad expression, translational fusions also revealed ZP protein production within multiple cell types that did not have that protein in their aECM (Figure 2C,D). In the case of LET-653(ZP) transgenes, expression and protein accumulation were detected only within 2° cells despite matrix assembly on 1° cells (Figure 2C). In the case of NOAH-1, endogenous protein was detected within all 2° cells (Figure 2D) despite surface enrichment only on vulC (Figure 1D). Therefore, ZP matrix assembly does not match sites of ZP gene transcription or protein expression and must depend on some other feature of the destination cell type.

### Lineal origins and ZP matrix assembly are separable components of vulva cell identity

Signaling by LET-23/EGFR and LIN-12/Notch specifies VPC fates (Shin and Reiner, 2018) (Figure 1B, 3A). Whether a VPC adopts the 1° or 2° fate is traditionally assessed by the pattern of divisions it undergoes after induction (lineage cell type) (Sternberg and Horvitz, 1986) and/or by properties of its descendants such as expression of specific marker transgenes (Gupta et al., 2012; Inoue et al., 2005). A limitation in the field has been the small number of cell identity markers available. Our data indicate that ZP matrix assembly is another marker of cell identity, so we focused on these three ZP proteins to investigate how the matrix on 1° cells is specified as distinct from the matrix on the 2° cells. Proper 1° matrix should include LET-653(ZP) and CUTL-18 and exclude NOAH-1.

**Figure 3:**
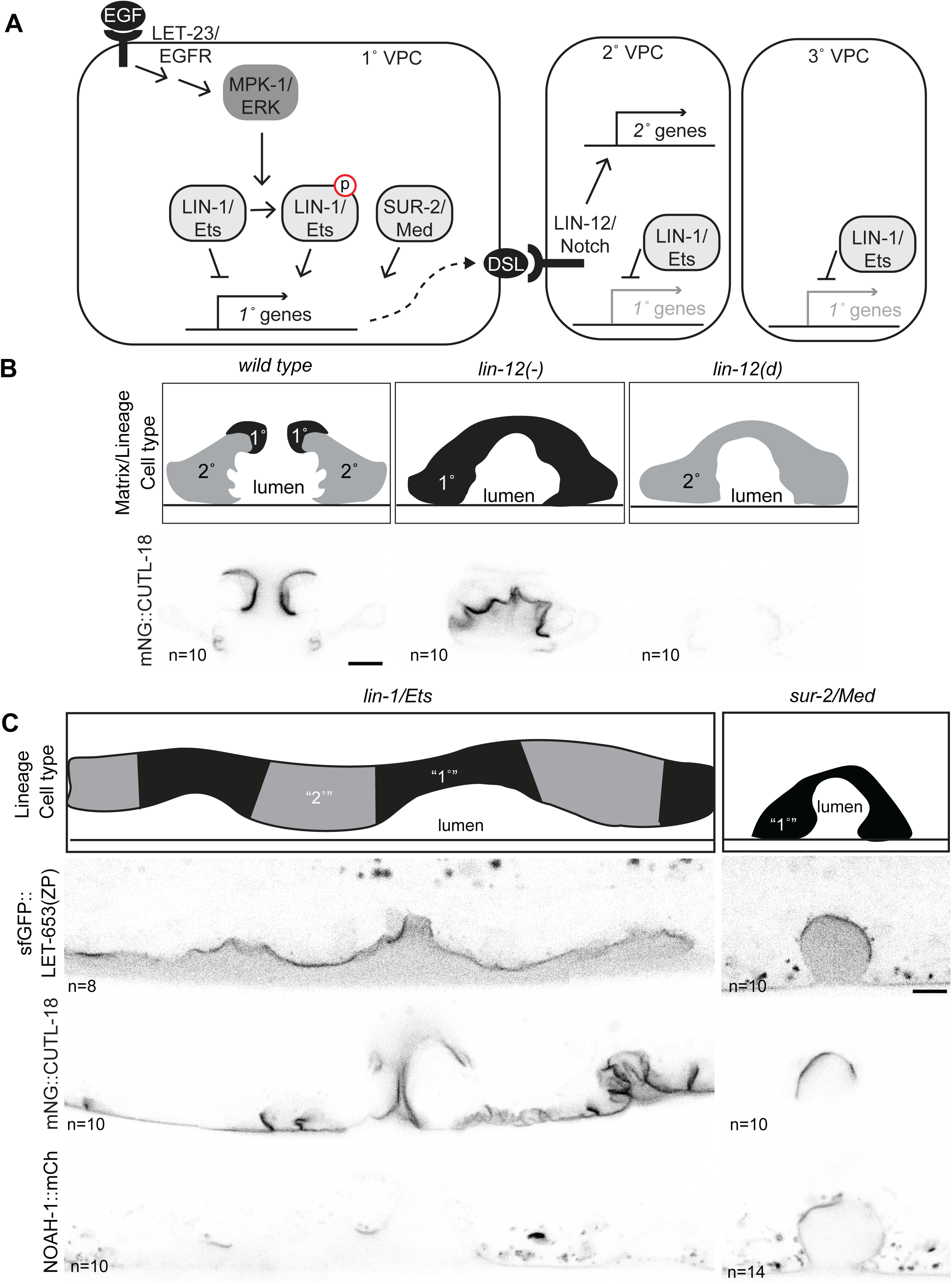
Contributions of EGFR effectors to matrix identity A) Transcription factors downstream of MPK-1/ERK promote transcription of 1° expressed genes. In the 1° VPC, LET-23/EGFR activates MPK-1/ERK via a Ras signaling cascade. MPK-1 phosphorylates LIN-1/ETS to convert it from a repressor to an activator of 1° identity (Jacobs et al., 1998; Tan et al., 1998). The mediator component SUR-2 acts in parallel to LIN-1 to promote some aspects of 1° identity, such as expression of DSL family ligands that activate LIN-12/Notch to promote the 2° fate in neighboring VPCs (Singh and Han, 1995; Underwood et al., 2017; Zhang and Greenwald, 2011). Unphosphorylated LIN-1 represses 1° expressed genes in the 2° and 3° lineages. B) Matrix identity *lin-12*/Notch mutants aligns with identity assessed by lineage. Top: schematic of vulva cell types (as assessed by lineage or matrix) in wild type, loss of function (-) *lin-12*(*n137n720)* mutants and dominant gain of function (*d*) *lin-12(n137)* mutants (Greenwald et al., 1983). Bottom: CUTL-18 is strongly enriched on all cell surfaces *lin-12(-)* mutants, consistent with a 1° matrix identity, but at faintly on some cell surfaces in *lin-12(d)* consistent with a 2° matrix identity. Medial confocal slices of mid-L4 stage worms. C) LIN-1/Ets represses, SUR-2/Med promotes 1° matrix identity. Top: schematic of vulva cell fates determined by lineage *lin-1(n304)* and *sur-2(ku9)* mutants (Beitel et al., 1995; Howard and Sundaram, 2002; Singh and Han, 1995). Below: LET-653(ZP)::sfGFP (*csIs66)* and CUTL-18 are enriched on the surface of both 1° lineage (orange arrowhead) and 2° lineage (blue arrow) vulva cells in a *lin-1* mutant. Few cells have NOAH-1::mCh on their surface in *lin-1* mutants. LET-653(ZP)::sfGFP, CUTL-18 and NOAH-1 are all enriched on a subset of 1° lineage cells in *sur-2* mutants. All scale bars 5µm.

In the absence of *lin-12/Notch,* all induced VPCs adopt a 1° fate as assessed by lineage. By the same metric, gain of function *lin-12/Notch* mutations instead cause all VPCs to adopt a 2° fate (Greenwald et al., 1983). Previously we reported that a *lin-12(-)* vulva (only 1° cells) had LET-653(ZP) on all cell surfaces while a *lin-12(d)* vulva (only 2° cells) excluded LET-653(ZP) from all cell surfaces (Cohen et al., 2020b). The 2° matrix protein NOAH-1 had the opposite behavior. We observed the localization of CUTL-18 in the same *lin-12* mutants. Consistent with its wild-type pattern of strong enrichment on 1° descendants vulE/F and weak enrichment on 2° descendants vulB1/B2, CUTL-18 was strongly enriched on the surfaces of *lin-12(-)* vulvas and weakly enriched on the ventral-most cells of *lin-12(d)* vulvas (Figure 3B). Thus, when cell fate specification is disrupted by *lin-12*/Notch mutations, the matrix and lineage cell type are in agreement.

However, we found that this relationship breaks down in mutants for the EGFR pathway effectors LIN-1 and SUR-2, which have more complex effects on VPC identity. LIN-1 is an Ets domain transcription factor and key substrate of MPK-1/ERK that both inhibits and promotes different aspects of 1° VPC identity (Figure 3A) (Beitel et al., 1995; Jacobs et al., 1998; Tuck and Greenwald, 1995). *lin-1* mutants are Muv with alternating 1° and 2° lineages as assessed by cell divisions (Beitel et al., 1995). Nevertheless, we observed that many more *lin-1* vulva cells than expected behaved like vulE/F and assembled LET-653(ZP) and CUTL-18 on their surface, whereas few cells recruited NOAH-1 (Figure 3C). Therefore, the matrix identity of *lin-1* vulva cells is distinct from the fate assessed by lineage, with some 2° lineage descendants having a 1°-like matrix identity (Fig. 3C). An endogenous LIN-1::GFP reporter (Kudron et al., 2018) was expressed in all VPCs and their immediate descendants, but expression then decreased and was barely detectable in the vulva by mid-L4 (Supplemental Figure 3A). We conclude that LIN-1 likely functions at an early step of lineage specification to repress 1° matrix identity (Figure 3C).

Another effector of the EGFR signaling cascade is the mediator subunit SUR-2/Med23, which promotes some aspects of 1° VPC identity, including the transcription of DSL genes to signal adjacent cells to adopt the 2° fate through LIN-12/NOTCH (Singh and Han, 1995; Underwood et al., 2017; Zhang and Greenwald, 2011)(Figure 3A). *sur-2* mutants are either completely vulvaless or have a single induced VPC that divides in the 1° pattern, resulting in 8 vulva cells (Singh and Han, 1995). Nevertheless, only some of the vulva cells in *sur-2* mutants had LET-653(ZP) or CUTL-18 on their surface, while some had NOAH-1 on their surface, suggesting that mutant cells have a hybrid identity (Figure 3C). We conclude that SUR-2 promotes 1° vs. 2° matrix identity, likely by acting with unknown transcription factors (Figure 3A), and that matrix assembly is separable from lineage division pattern and other aspects of cell identity.

### LIN-11/LIM and COG-1/Nkx are not required for 1° vulva cell matrix assembly

Transcription factors LIN-11 and COG-1 are expressed in all vulva cells (Palmer et al., 2002) (Supplementary Figure 3B). Though *lin-11* mutants originally were identified based on defects in 2° cell fate (Ferguson et al., 1987), both LIN-11 and COG-1 are required for different aspects of both 1° and 2° cell fate execution (Gupta et al., 2003; Palmer et al., 2002). We tested whether either was required for LET-653(ZP) localization to 1° matrix. Although lumen shape was greatly disrupted, both *lin-11 null* and *cog-1(sy275)* missense mutants had LET-653(ZP) enriched on vulva cell surfaces at the apex of the vulva (Supplementary Figure 3C). Although the *cog-1* allele is not null, it affects a critical rescue within the DNA binding domain and has been shown to disrupt 1° fate execution (Palmer et al., 2002). This suggests that LIN-11 and COG-1 are not required for 1° matrix identity.

### EGL-38/Pax expression correlates with 1° vulva cell matrix assembly

The paired box transcription factor EGL-38 is expressed in the vulD, E and F cells and is required for vulE/F lumen shape but not all aspects of 1° fate execution (Mok et al., 2015; Webb Chasser et al., 2019) (Figure 4B). We assessed the expression of EGL-38::GFP in *lin-12/Notch* and EGFR effector mutant vulvas. Consistent with it being a robust marker of the 1° lineage and vulE/F identity, EGL-38::GFP was present in all vulva nuclei of *lin-12(-)* mutants (Figure 4A, B,E). Furthermore, most vulva cells in *lin-1* mutants expressed EGL-38::GFP (Figure 4C,D,E). Only some of the vulva cells in *sur-2* mutants expressed EGL-38::GFP (Figure 4C,D,E). We conclude that LIN-1 represses, and SUR-2 promotes EGL-38 expression and that EGL-38 expression correlates well with 1° matrix identity (Figure 4F).

**Figure 4:**
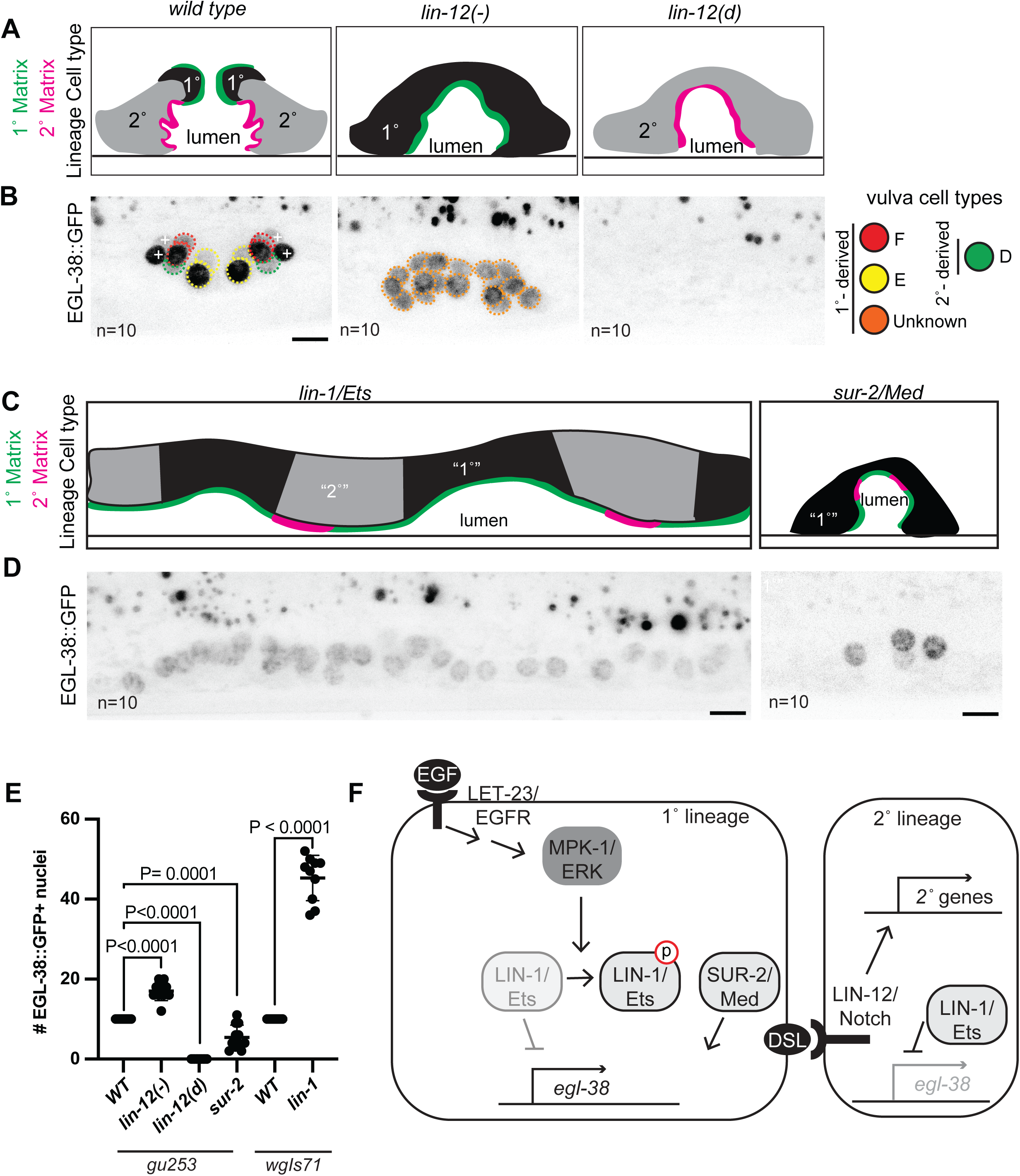
Vulva matrix identity correlates with EGL-38 expression A) Schematic of vulva cell types (as assessed by lineage) and summary of matrix assembly as in Figure 3 and (Cohen et al., 2020b) in *lin-12*/Notch mutants. B) EGL-38 expression in *lin-12*/Notch mutants aligns with matrix identity. Maximum projections of confocal Z stacks through entire vulva of mid-L4 stage worms. Vulva nuclei expressing EGL-38::GFP are circled - vulD in green, vulE in yellow, vulF in red, and unknown nuclei descended from a 1°-like lineage in orange. + indicates uv1 nuclei. n= number of worms, number of EGL-38 positive vulva nuclei per worm quantified in C. C) Schematic of vulva cell types (as assessed by lineage) and summary of matrix assembly as in Figure 3 in *lin-1* and *sur-2* mutants. D) LIN-1/Ets represses, SUR-2/Med promotes *egl-38* expression consistent with matrix identity. EGL-38::GFP expression in the indicated mutants. Maximum projections of confocal Z stacks through entire vulva of mid-L4 stage worms. Consistent with their matrix identity, the majority of vulva cells expressed EGL-38 in *lin-1* mutants, and only some of the vulva cells expressed EGL-38 in *sur-2* mutants. n= number of worms, number of EGL-38 positive vulva nuclei per worm quantified in E. All scale bars 5µm. E) Count of the number of EGL-38-positive vulva nuclei in wild type (WT) and *lin-12*, *lin-1,* and *sur-2* mutants. Presence of EGL-38 was assessed with EGL-38::GFP expressed from the endogenous locus (*gu253)* or EGL-38::TY1::EGFP::3xFLAG expressed from a transgene *(wgIs171)*. In wild type worms, EGL-38 was always present in 10 vulva nuclei, regardless of method of expression. P values Kruskal–Wallis test. F) Contributions of EGFR effectors to *egl-38* expression. LIN-1 represses *egl-38* transcription in the absence of EGFR/MAPK signaling. SUR-2 promotes *egl-38* transcription in the presence of EGFR/MAPK signaling.

Surprisingly, EGL-38 was not present in any vulva nuclei in *lin-12(d)* mutants, though it was present in the 2° lineage descendant vulD in wild type (Figure 4B,E). This suggests that EGL-38 expression in vulD may depend on the specification of the 1° lineage.

### EGL-38/Pax promotes 1° vulva cell matrix assembly

We next tested whether *egl-38* is required for 1° cell matrix gene expression. An *egl-38* null mutation is lethal shortly after hatch (Chamberlin et al., 1997; Clark, 1990). Two missense mutations in the DNA-binding paired box domain (*sy294* and *n578*) have a normal 1° cell lineage division pattern (Trent et al., 1983), but cause vulva morphology defects (Figure 4A) (Chamberlin et al., 1997; Rajakumar and Chamberlin, 2007). We focused on the allele *(n578)* with higher penetrance of the vulva defect. In *egl-38(n578)* mutants, the *let-653* and *noah-1* transcriptional reporters were still expressed in all vulva cells (Supplementary Figure 4), but the *cutl-18* reporter showed reduced expression specifically in vulE/F (Figure 5C,D). Therefore, EGL-38 does promote *cutl-18* expression as one aspect of vulE/F matrix identity execution.

**Figure 5:**
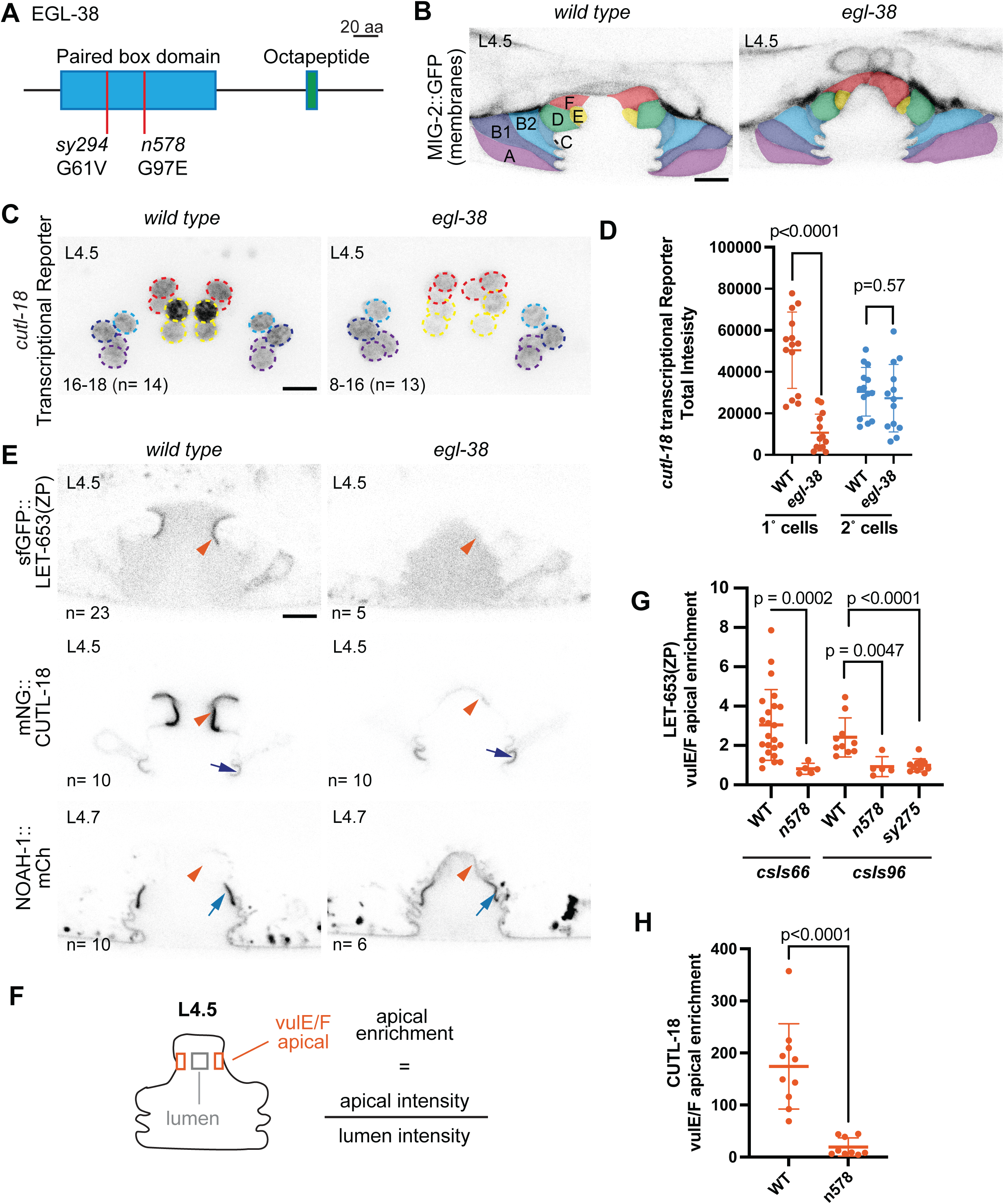
The Pax transcription factor EGL-38 is required for proper 1° cell ZP matrix assembly A) Schematic of EGL-38 protein. Red bars indicate the location of hypomorphic missense mutations *n578* and *sy294* in the conserved Paired box (Pax) domain. B) In *egl-38* mutants the vulF cells remain close to each other, blocking the uterine connection. Vulva cells visualized with the membrane marker MIG-2::GFP(*muIs28*) and colored by cell type. C) EGL-38 promotes the transcription of *cutl-18* in 1° descendants vulE/F. Maximum projections of confocal Z stacks through entire vulva of mid-L4 of wild type and *egl-38(n578)* worms expressing the indicated transcriptional reporter. Vulva nuclei are outlined according to the cell colors in A. Lower left: Number of mCherry positive nuclei, (n= number of worms). Wild type Ns represent the same worms as in Figure 2. *let-653* and *noah-1* transcription were not affected, see Supplemental figure 4. D) Quantification of *cutl-18* transcriptional reporter fluorescence in 1°- and 2°-descendant nuclei in wild type and *egl-38(n578)* mutants. See methods. P values Kruskal–Wallis test. E) EGL-38 is required for proper 1° cell ZP matrix assembly. LET-653(ZP)*csIs66*, mNG::CUTL-18 and NOAH-1::mCherry in wild-type and *egl-38(n578)* mutants. In *egl-38* mutants the 1° matrix (orange arrowheads) does not contain the proper proteins, while 2° cell specific matrices (blue arrows) are maintained. Medial confocal slices at the indicated L4 stages. (n= number of worms). Wild type Ns represent the same worms as in Figure 1. See Supplemental Figure 5 for images of additional alleles and fluorescence quantification of 2° cell surfaces. All scale bars 5 µm. F) Apical enrichment of ZP proteins quantified by fluorescence intensity at the vulE/F surfaces divided by intensity in a box of the same total size in the lumen. See Methods. G and H) Apical enrichment of LET-653(ZP) or CUTL-18 on vulE/F cells. Column labels below indicate wild type worms (WT) or *egl-38* alleles (*n578* and *sy294)* and LET-653(ZP) transgenes (*csIs66* and cs*Is96).* All P values Kruskal–Wallis test.

We observed the localization of tagged ZP proteins in wild type and *egl-38* mutant vulvas and quantified their enrichment on apical cell surfaces (Figure 5E-H, Supplementary Figure 5A-F). In *egl-38(n578)* mutants, both LET-653(ZP) and CUTL-18 showed severely reduced enrichment on 1° cells (Figure 5E, G, H), while NOAH-1 showed aberrant recruitment to 1° cells (Figure 5E, Supplementary Figure 5E). This pattern was independent of the LET-653(ZP) transgene tested (Figure 5G, Supplementary Figure 5A). CUTL-18 and NOAH-1 still localized to specific 2° cell surfaces in *egl-38* mutant vulvas; however, their level of enrichment changed from wild type, suggesting there may be a finite pool of protein that can move between cell surfaces (Figure 5E, Supplementary Figure 5C, F). We tested whether the absence of proper 1° matrix was linked to the failure of vulF cells to separate in *egl-38* mutants. While the majority of *egl-38(sy294)* worms have normal vulva morphology (Rajakumar and Chamberlin, 2007), we found these mutants have the same LET-653 and NOAH-1 localization defects as in *egl-38(n578)* (Figure 5G, Supplementary Figure 5A, E, F). We conclude that *egl-38* is required to promote 1° vs. 2° matrix assembly and that its targets are likely to include factors that recruit or modify specific ZP proteins.

## Discussion

All epithelial cells are covered by aECMs that play important roles in tissue shaping and barrier functions, but how cells assemble the proper type of aECM is still poorly understood. Here we investigated cell-type specific aECM as a feature of epithelial cell identity in the *C. elegans* vulva, where cell fates are controlled by a combination of EGFR and Notch signaling and a network of downstream transcription factors. Different ZP domain proteins assemble on the surfaces of different vulva cell types, but these patterns do not reflect cell-type specific expression of the ZP proteins themselves. Rather, ZP proteins are broadly expressed, and unknown features of the destination cell types allow each to recruit and assemble the appropriate set of ZP proteins on their surface. LET-23/EGFR-dependent 1° vulva cell types express the Pax2/5/8 family transcription factor EGL-38, which promotes 1° matrix assembly (Figure 6) but is not required for other aspects of 1° fate execution. Our results connect the known signaling pathways and various downstream effectors to EGL-38/Pax expression and the ZP matrix component of vulva cell fate execution. We propose that different epithelial subtypes may use transcription factors like EGL-38 to turn on batteries of genes controlling their unique aECM biology.

**Figure 6:**
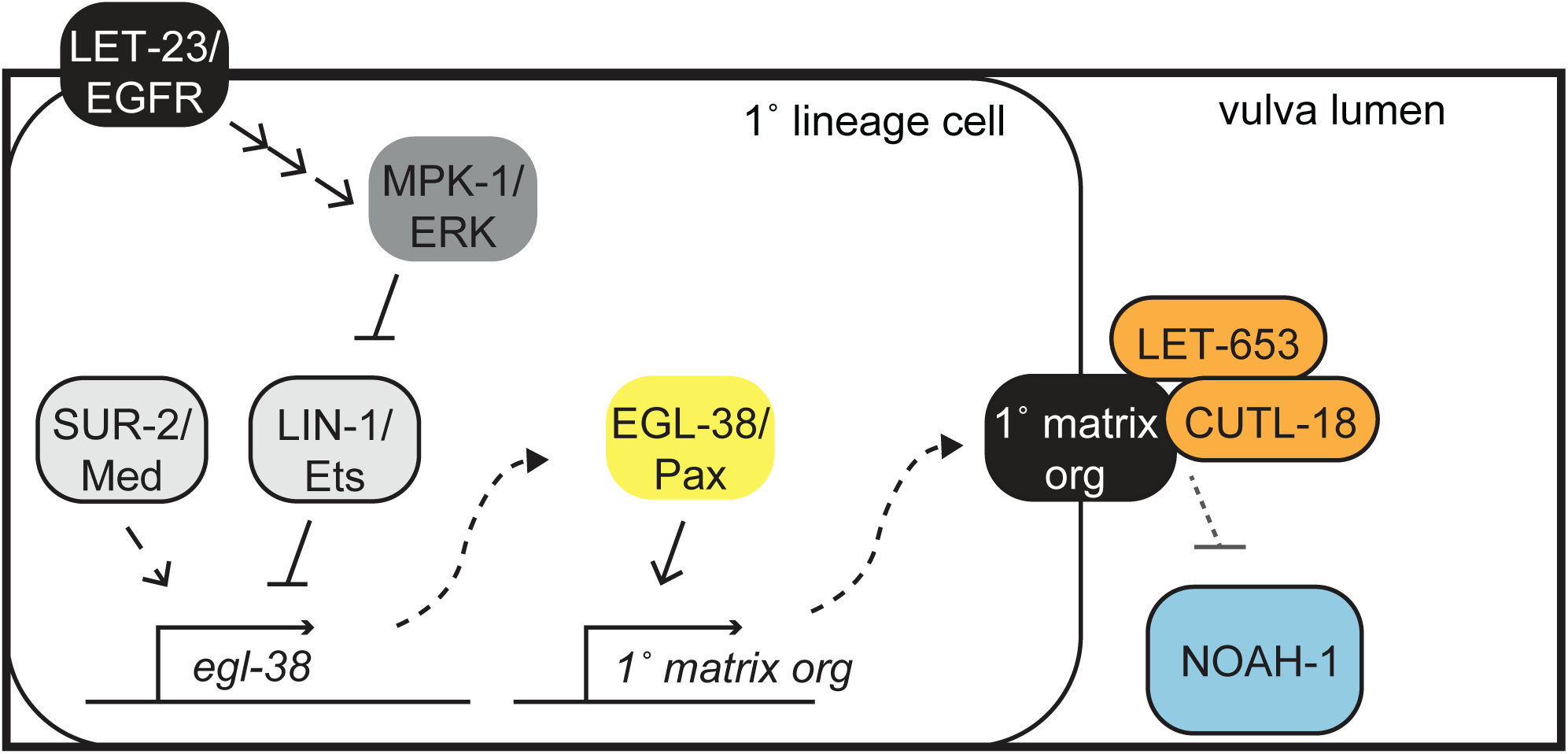
Model for 1° matrix fate execution The 1° lineage is specified through LET-23/EGFR, triggering a signaling cascade to MPK-1/ERK. Signaling promotes *egl-38* expression through at least two transcription factors, by relieving LIN-1/Ets repression through direct phosphorylation and promoting SUR-2/Med activity (or that of an unknown partner transcription factor). EGL-38 in turn promotes the expression of unknown 1° matrix organizer(s) that are present on the surface of primary cells and recruit 1° matrix proteins LET-653(ZP) and CUTL-18 and exclude 2° matrix protein NOAH-1.

### How are ZP proteins recruited to specific cell surfaces?

ZP domain proteins are commonly produced in and localize to the surface of different cell types, sometimes separated by large distances. For example, some ZP protein components of chicken and fish egg coats are produced in the liver and then travel through the bloodstream to reach the oocytes (Bausek et al., 2000; Conner and Hughes, 2003). The mammalian intestinal ZP protein GP2 is produced primarily in the pancreas (Kurashima et al., 2021), while alpha-tectorin is widely expressed in the sensory, roof, and transitional epithelia of the cochlear but assembles into specific patterns in the tectorial membrane (Kim et al., 2019; Niazi et al., 2024; Rau et al., 1999). Here we found that three ZP proteins were expressed broadly but localized to specific cell surfaces in the vulva. How ZP proteins are properly targeted to their destination cell surface remains an open question.

One contributing factor to ZP protein assembly is cleavage at a site immediately following the ZP domain. Many ZP proteins are cleaved at a consensus furin cleavage site (RxxR) (Bokhove and Jovine, 2018), although some can also be cleaved at non-RxxR sites by other types of serine proteases (Brunati et al., 2015; Niazi et al., 2024). We previously showed that the RxxR cleavage site in LET-653(ZP) was required for its assembly in the vulE/F matrix (Cohen et al., 2020a); however, the relevant protease(s) acting on this site remain unknown. This cleavage is predicted to relieve an intramolecular interaction between the ZP domain and the protein C-terminus, allowing the ZP domain to bind to other proteins and assemble in the matrix (Jovine et al., 2004). Thus, spatial control of relevant proteases could control spatial assembly of ZP matrix proteins.

Another contributing factor to ZP protein assembly may be the presence of relevant binding partners. Some ZP proteins, including LET-653, have no membrane associated domain (Figure 1C). Others, including NOAH-1 and CUTL-18, have transmembrane domains or GPI anchors that are predicted to be cleaved from the ZP domain (Figure 1C). Consequently, mature ZP proteins likely bind to a membrane-associated partner in their target matrix. In the case of LET-653, the C-terminal half of the ZP domain is sufficient to confer 1° matrix localization, suggesting this domain must bind a relevant partner (Cohen et al., 2020a). Some ZP domains are known to bind other ZP proteins or collagens, which can affect their localization (Andrade et al., 2016; Chu and Hayashi, 2021; Ghosh and Treisman, 2024; Niazi et al., 2024). While we found CUTL-18 and LET-653 are not required for each other’s localization to the vulE/F matrix, *C. elegans* has 43 ZP proteins and 173 cuticle collagens (Cohen et al., 2019; Teuscher et al., 2019; Weadick, 2020), some of which could be binding partners. Alternatively, ZP proteins could bind other types of transmembrane proteins (Hölzl et al., 2011; Pfistershammer et al., 2008). Relevant candidates could be further narrowed by identifying those with vulva cell-type specific expression patterns. Overall, the vulva has a small number of cells, yet multiple cell-type specific ZP matrices, making it an ideal model for further study of how ZP proteins localize to the proper matrix.

### What is the purpose of organizing the aECM into cell-type specific domains?

The large number of *C. elegans* ZP proteins, coupled with their cell type specific matrix incorporation (Figure 1)(Cohen et al., 2020a, 2019; Yu et al., 2000), suggests different ZP proteins have different matrix functions. For example, different ZP proteins may bind and recruit other matrix components or confer different physical properties that affect tissue shaping. While we focused here on matrix within the developing vulva, some ZP proteins (such as CUTL-18) remain as part of the adult cuticle, and even transient pre-cuticle proteins can influence the structure of the subsequent cuticle (Cohen et al., 2019; Katz et al., 2022; Mancuso et al., 2012; Serra et al., 2024). In the adult vulva, the apical surfaces of the 1° descendants remain internal, while the 2° descendants are oriented outwards. Some vulva cells attach to the uterine or vulva muscles which contract to drive egg-laying. These conditions could require differences in rigidity. Components of the cuticle can also affect pathogen resistance and mate attraction (Gravato-Nobre and Hodgkin, 2005; Weng et al., 2023). *let-653* mutants have relatively mild defects in late stages of vulva morphogenesis (Cohen et al., 2020b) and we did not observe any obvious changes in *cutl-18* mutants (Supplementary Figure 2), but it will be important to test for ultrastructural, physiological and behavioral changes to understand the contributions made by the specific matrices on different vulva cell types.

### How does EGL-38/PAX promote 1° matrix assembly and influence vulva lumen shape?

Pax family transcription factors can be “master regulators” of cell identity or they can regulate morphogenesis (Shaw et al., 2024; Thompson et al., 2021). EGL-38 appears to promote only a subset of vulE/F properties, including matrix assembly, rather than being a master regulator of vulE/F identity. One caveat is that the *egl-38* alleles studied are hypomorphs since the null is lethal. It is possible that EGL-38 may have a larger role in vulE/F identity.

We propose that EGL-38 promotes the assembly of 1° matrix by promoting the expression of one or more unknown “1° matrix organizers” that contribute to ZP protein assembly, such as through proteolysis or binding. Identification of such matrix organizers has been challenging given the few known EGL-38 targets and limited information about preferred EGL-38 binding motifs (Johnson et al., 2001; Webb Chasser et al., 2019). Although EGL-38 is a transcriptional activator of its few known targets (Johnson et al., 2001; Webb Chasser et al., 2019), it could also act as a repressor on some targets, as many Pax transcription factors act as both (Shaw et al., 2024). Transcriptomic studies are currently underway to identify *egl-38*-dependent genes in the vulva.

Given the known roles of aECM in tube lumen shaping (Li Zheng et al., 2020; Sundaram and Pujol, 2024), defects in 1° matrix assembly could potentially explain the failure of the vulF cells to separate properly in *egl-38* mutants. However, although we found EGL-38 promoted *cutl-18* expression in vulE/F and the assembly of both CUTL-18 and LET-653 in the matrix, these changes alone are not sufficient to explain the phenotype since vulF still can separate when these ZP proteins do not assemble (Supplementary Figure 5A). *egl-38* mutants must have additional changes in the 1° matrix or in other relevant cellular processes, such as anchor cell invasion (Estes and Hanna-Rose, 2009; Kenny-Ganzert and Sherwood, 2024), that remain to be discovered.

### The vulva as a model for connecting transcription factor networks to cell and matrix biology

The seven vulva cell types differ in many properties, including cell shape, dorsal vs. ventral adhesion, cytoskeletal organization, and matrix organization (Cohen et al., 2020b; Gupta et al., 2012; Mok et al., 2015; Sharma-Kishore et al., 1999). The transcriptional logic that specifies these various differences is just beginning to be deciphered (Farooqui et al., 2012; Fergin et al., 2022; Sherwood and Plastino, 2018; Underwood et al., 2017). Our data suggest that in 1° vulva cell types, LET-23/EGFR signaling promotes EGL-38/Pax expression by simultaneously relieving LIN-1/Ets-dependent repression and promoting SUR-2/Med23-dependent transcription (Figure 6). EGL-38 may then promote expression of gene batteries related to a subset of cell properties, such as matrix organization, while other transcription factors such as LIN-11 and COG-1 promote other properties. EGL-38 may also function cooperatively with other transcription factors to promote additional aspects of vulE/F biology (Fernandes and Sternberg, 2007). Notably, EGL-38 is expressed only transiently and disappears from vulva cells by the late L4 stage, so it does not fit the classical definition of a “terminal selector” that functions continuously to maintain unique aspects of cell identity (Hobert and Kratsios, 2019). Instead, the period of EGL-38 expression coincides well with vulva morphogenesis and the times when different aECMs are built and remodeled. EGL-38-dependent targets may be less relevant in the adult, when distinct transcriptional networks likely take over to maintain cell identity and appropriate physiological functions. Going forward, we expect that single cell transcriptomic data will reveal key differences in gene expression between vulva cell types and stages, help identify candidate matrix organizers and other features of each cell type’s unique biology, and allow dissection of the transcriptional networks that establish these differences. Just as the *C. elegans* vulva has served as a paradigm for signal-dependent patterning of cell fates (Shin and Reiner, 2018), it can also provide a useful model for understanding finer aspects of epithelial identity control that may be broadly relevant to other epithelia across organisms.

## Methods

### Worm strains and maintenance

All *C. elegans* were cultured at 20°C, according to standard methods (Brenner, 1974). The tagged ZP domain of LET-653 (LET-653(ZP)) was expressed from either of two transgenes *csIs66*[*let-653pro::sfGFP::LET-653(ZP); let-653pro::PH::mCherry*] or *csIs96*[*let-653pro::LET-653(ZP)::sfGFP; lin-48pro::mRFP*], which have identical localization and rescue a *let-653(cs178)* null (Cohen et al., 2020a). Novel alleles generated by CRISPR/Cas9 genome editing (*cutl-18(syb8315), cutl-18 (syb8442), let-653(syb8336),* and *noah-1(syb8346)*) were designed using genome and transcript sequences from Wormbase with protein domain predictions from SMART and generated by Suny Biotech (Davis et al., 2022; Letunic et al., 2021). Strains used are listed in Supplementary Table 1.

### Microscopy

L4 substages were identified by size and vulva lumen morphology (Mok et al., 2015). Early adults were selected by picking L4 larvae and imaging those worms the following day. For confocal microcopy worms were immobilized with 10 mM levamisole in M9 buffer and mounted on 5% agarose pads supplemented with 2.5% sodium azide. Images were captured with a Leica TCS DMi8 confocal microscope through a 63x HC PL APO objective, Numerical Aperture 1.3, with a Z step size of 0.33µm using Leica LasX Software and processed in Fiji (Schindelin et al., 2012). Laser power and line accumulation settings varied with the tissue and tagged protein imaged but were held constant between wild type and mutant worms expressing the same tagged protein. The number of worms imaged is noted on the representative micrograph in each figure.

### *cutl-18* transcriptional reporter quantification

The total intensity of mCh::H2B fluorescence was measured inside a 171 µm^2^ rectangle containing the 1° cell nuclei and two 146 µm^2^ rectangles containing the 2° cell nuclei on each side of the vulva (total of 292 µm^2^), without background subtraction. Measurements were performed on sum projections of Z stacks through the entire vulva.

### Apical enrichment measurements

Measurements of ZP protein enrichment at vulva cell surfaces were performed based on the method previously described (Cohen et al., 2020a), modified to locate the apical surface in the absence of protein enrichment. Using the DIC channel of each image to locate the apical cell surface, the total intensity in the fluorescent channel was measured in a 10 pixel high by 5 pixel wide box on the left and right apical surfaces of the indicated vulva cell and a 10 pixel square in the vulva lumen. Apical enrichment was calculated as the sum of the two apical measurements divided by the lumen measurement (Figure 5F, Supplementary Figure 5B, D). All measurements are taken from single confocal Z-slices. Final values reported are ratios.

## Acknowledgements

We thank Michel Labouesse and Helen Chamberlin for strains, John Murray for helpful comments on the manuscript, and other members of the Penn Worm Group for thoughtful discussions. Some strains were obtained from the Caenorhabditis Genetics Center, which is supported by the National Institutes of Health Office of Research Infrastructure Programs P40 OD10440.

---

**Supplement 1:**
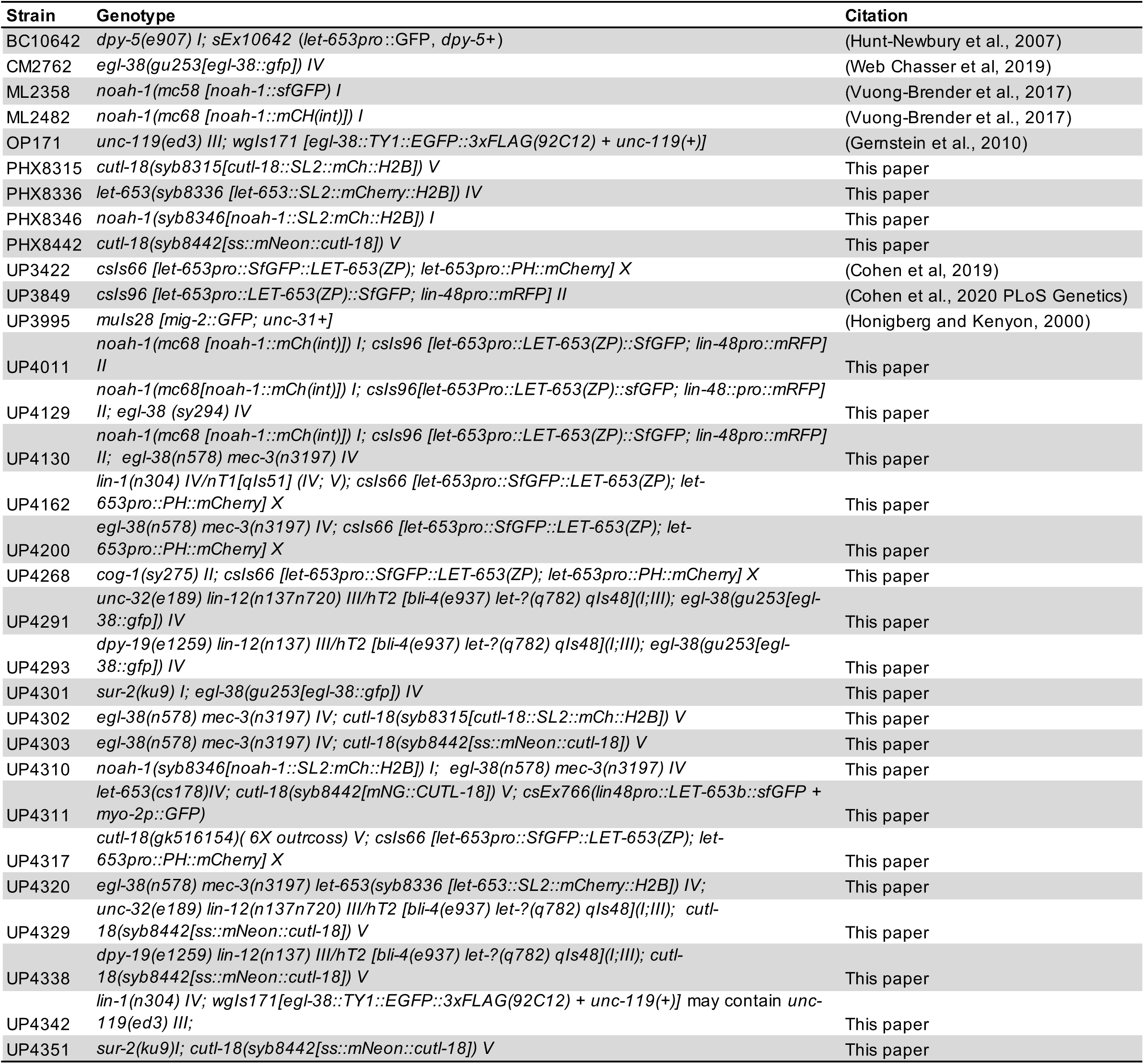
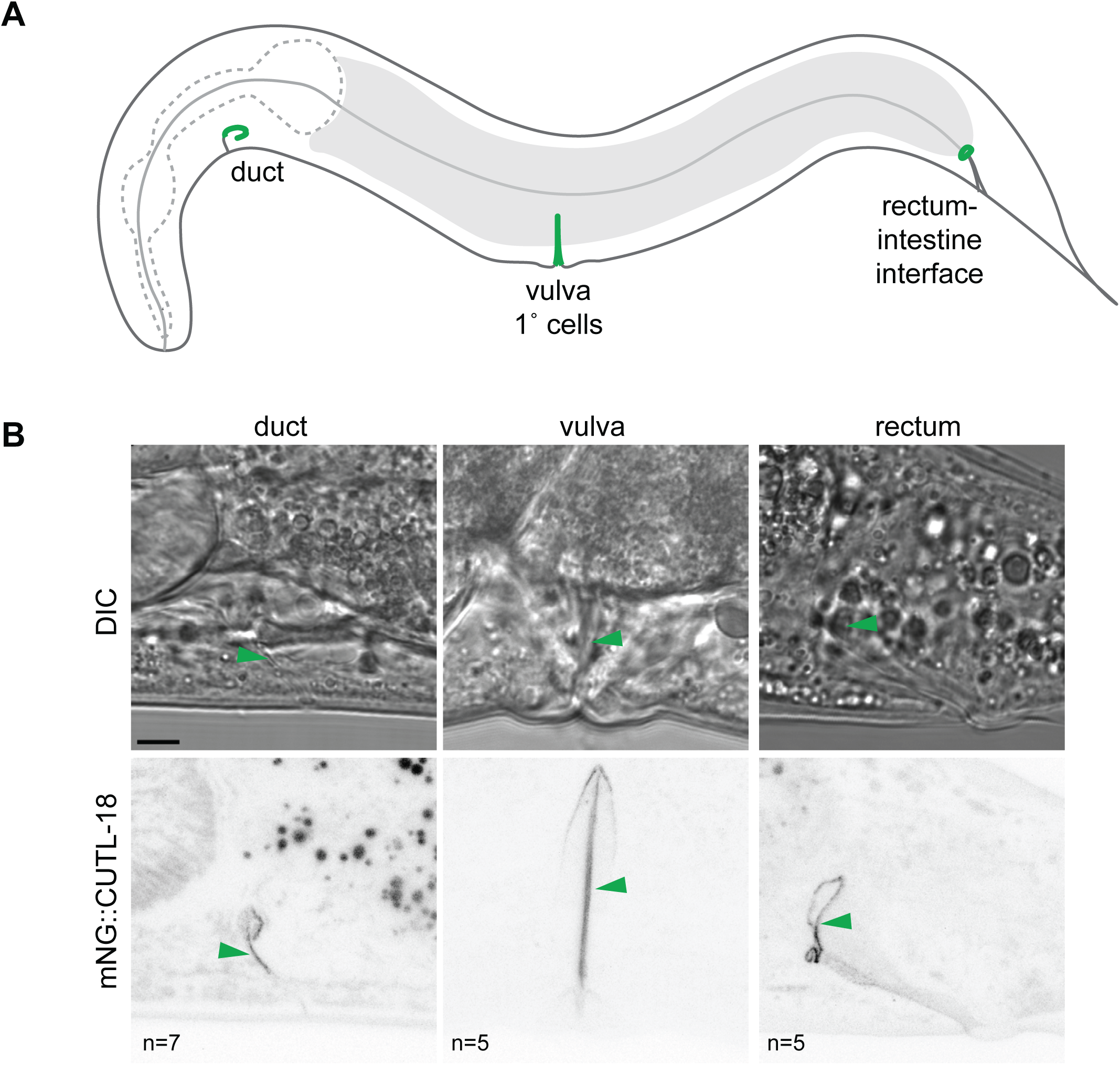
CUTL-18 is a cell-type specific cuticle component in multiple interfacial tubes A) Diagram of an adult worm. Green indicates the presence of mNG::CUTL-18. B) mNG::CUTL-18 localization in day 1 adult worms. Top row are DIC images of the indicated tissue. Bottom row are maximum projections of confocal Z stacks through the indicated tissue. Arrowheads point to location of mNG:CUTL-18 in both images. Scale bar 5 µm.

**Supplement 2:**
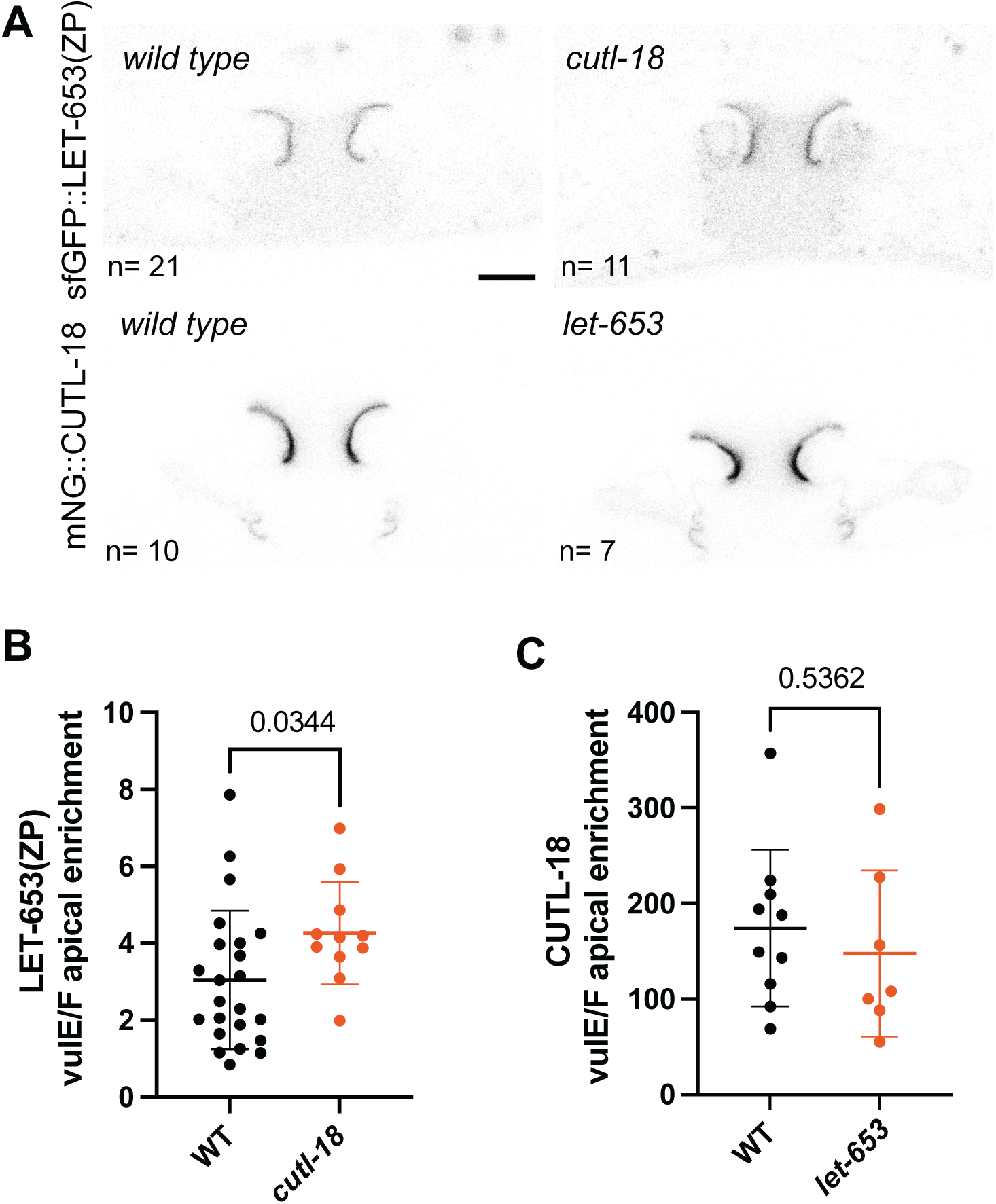
LET-653 and CUTL-18 do not depend on each other for 1° matrix assembly A) A) Mid-(L4.4/L4.5) L4 larval stage vulvas of wild type, *cutl-18(gk516154)* and *let-653(cs178)* worms expressing LET-653(ZP):sfGFP*(csIs66)* or mNG::CUTL-18. *cutl-18(gk516154)* is a A>T substitution in the splice acceptor site of exon 4 (Thompson et al., 2013), skipping exon 4 would result in a frameshift and premature termination. Outcrossed 6x. *let-653(cs178)* is a null mutation, rescued in the duct by *csEx766*(lin48pro::LET-653b::sfGFP + myo-2p::GFP) (Forman-Rubinsky et al., 2017). Medial confocal slices. Scale bar 5 µm. B) and C) Apical enrichment of LET-653(ZP) or CUTL-18 on vulE/F cells. See Figure 5F and Methods. P values Kruskal–Wallis test.

**Supplement 3:**
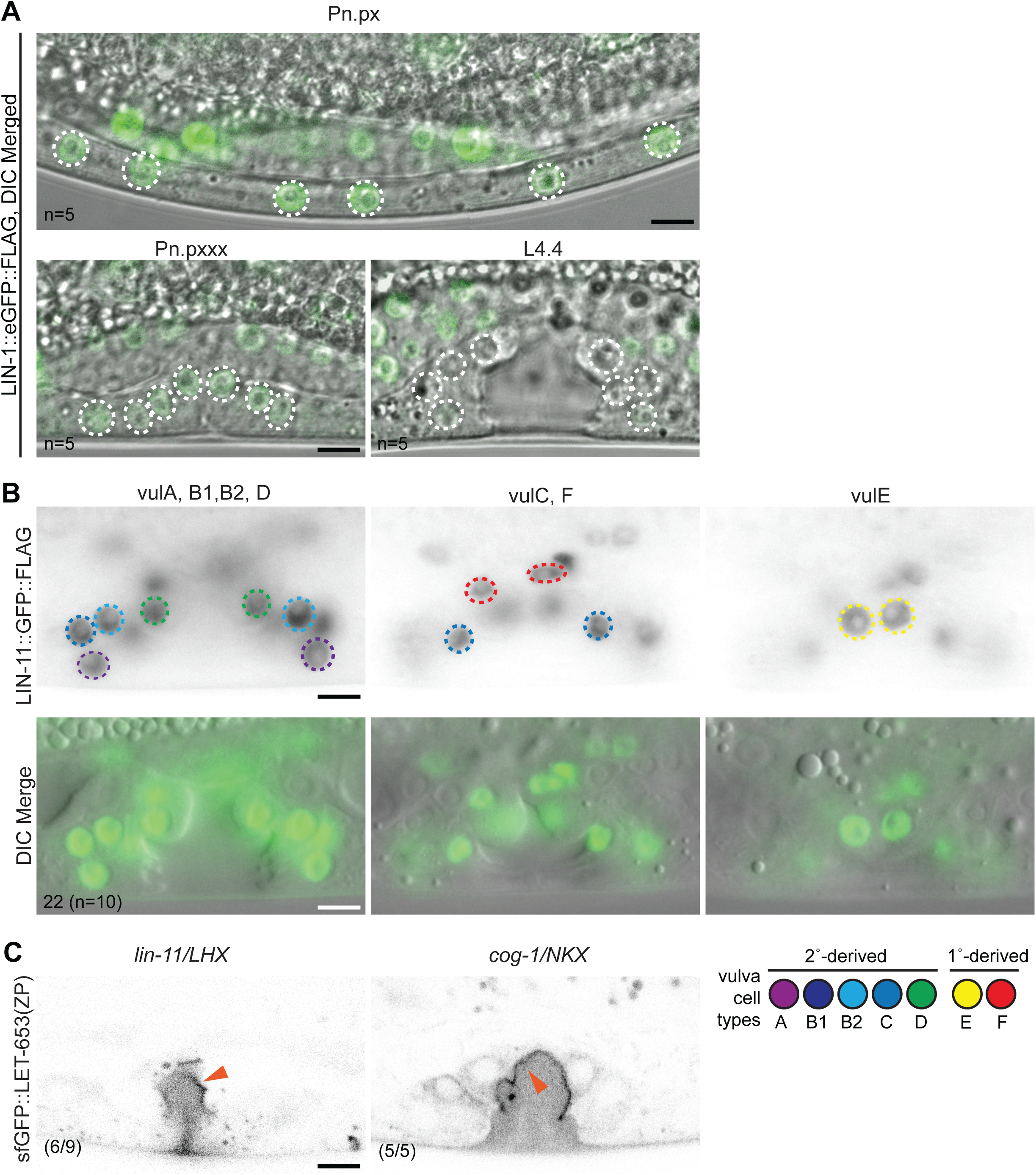
LET-653(ZP) assembly does not require other 1° expressed transcription factors. A) LIN-1::GFP is expressed in all VPC descendants during the L3 stage, but is not detectable in the vulva by L4. DIC + confocal merged images of the VPC descendants (circled) during L3 Pn.px and Pn.pxx, and L4.4 stage. B) LIN-11::GFP is expressed in all vulva cells. Inverted epifluorescence (top) and DIC + epifluorescence merged images (bottom) of the same L4.4 stage vulva at different focal planes with the above indicated vulva nuclei in focus. Lower left: number of nuclei expressing LIN-11::GFP (total number of mid-L4 (L4.4/L4.5) worms). C) LET-653(ZP)::sfGFP *csIs66* in mid-L4 vulvas of *lin-11(n389)* and *cog-1(sy295)* worms. LET-653(ZP) enriched on the surface of 1° cells (orange arrowhead) Lower left: N/total worms observed. Medial confocal slices at the indicated L4 stages. All scale bars 5 µm.

**Supplement 4:**
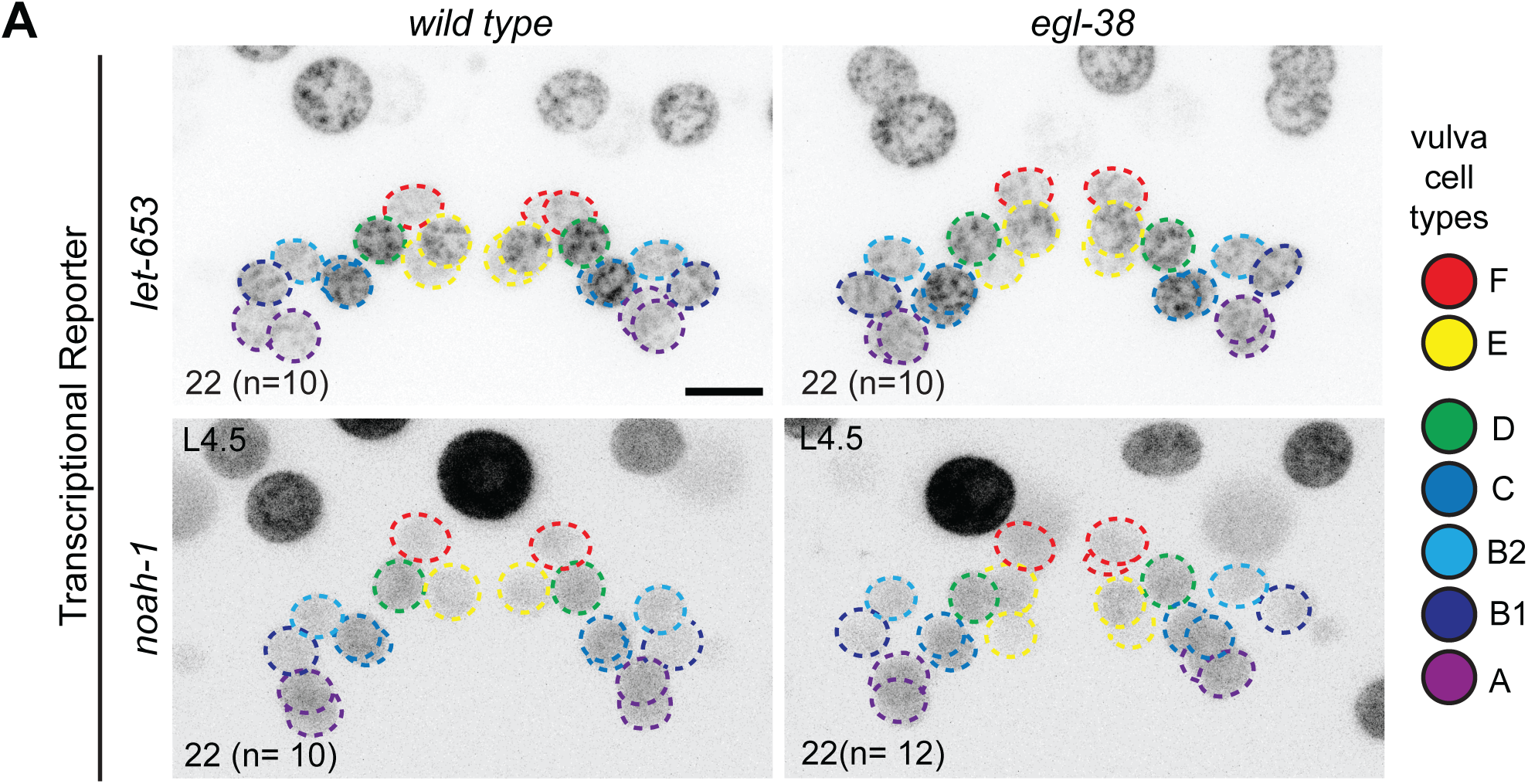
Transcription of *let-653* and *noah-1* does not require on *egl-38*. A) Maximum projections of confocal Z stacks through entire vulva of worms expressing the indicated transcriptional reporter. Vulva nuclei are outlined according to the adjacent color code. Scale bar 5 µm Lower left: Number of mCherry positive nuclei, (n= number of worms), wild type Ns represent the same worms as in Figure 2.

**Supplement 5:**
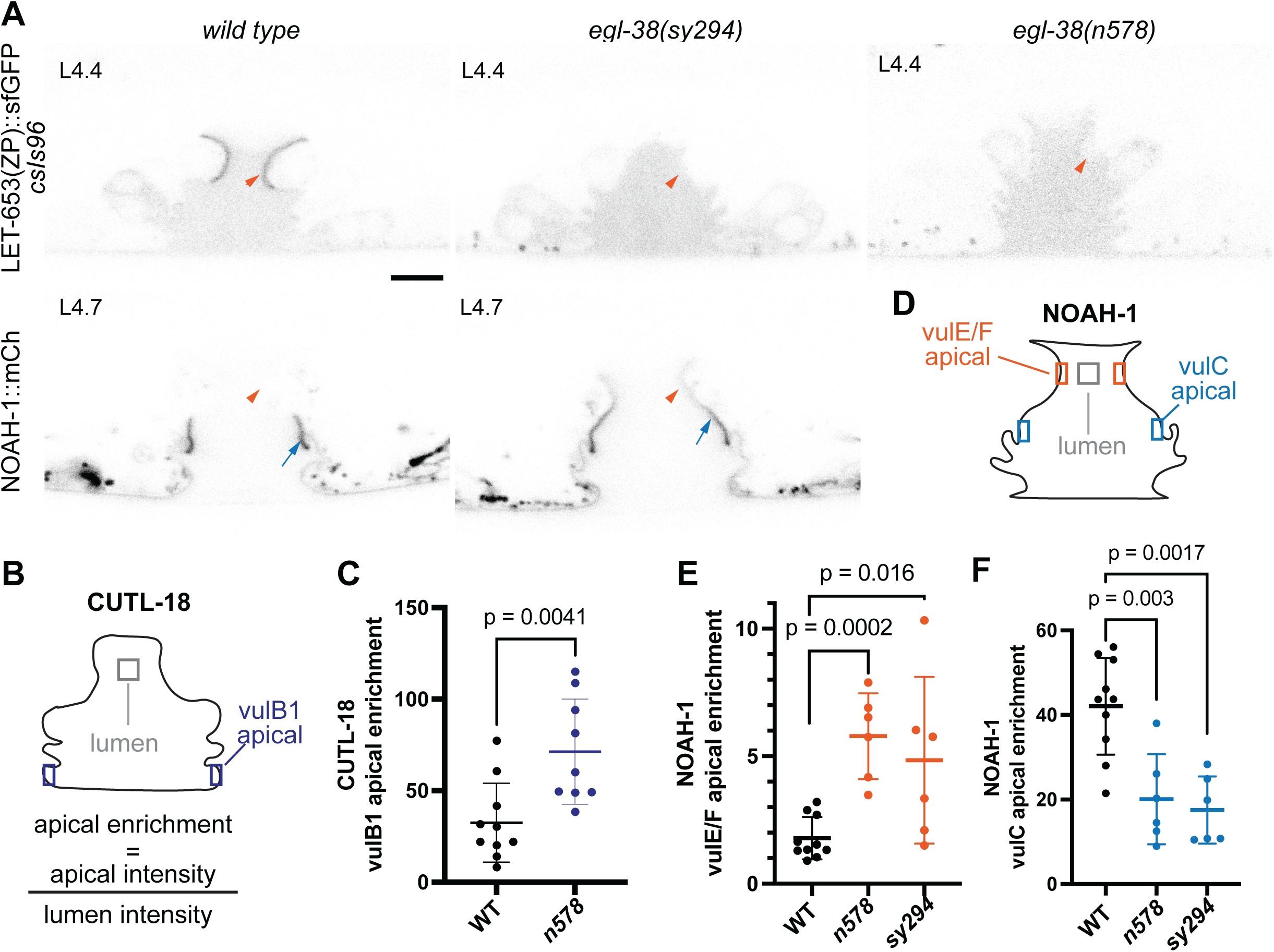
EGL-38 requirement for 1° matrix is independent of allele and *let-653(ZP)* transgene A) Vulvas of wild type and *egl-38* mutant worms at the indicated stage expressing LET-653(ZP)::sfGFP (*csIs96)* or NOAH-1:mCherry. As with LET-653(ZP)::sfGFP *csIs66* and *egl-38(n578)* (Figure 5E), in *egl-38* mutants the 1° matrix (orange arrowhead) does not contain the proper proteins, while 2° cell specific matrices (blue arrows) are maintained. Medial confocal slices at the indicated L4 stages. Scale bar 5 µm B) Apical enrichment of CUTL-18 quantified by fluorescence intensity at the vulB1 surfaces divided by intensity in a box of the same total size in the lumen. See Methods. C) Apical enrichment of CUTL-18 on vulB1. Column labels below indicate wild type worms (WT) or *egl-38* alleles (*n578*). D) Apical enrichment of NOAH-1 quantified by fluorescence intensity at the 2°descendant vulva cell surfaces, divided by intensity in a box of the same total size in the lumen. See Methods. E,F) Apical enrichment of NOAH-1 on the indicated the 2° descendant vulva cell surfaces Column labels below indicate wild type worms (WT) or *egl-38* alleles (*n578* and *sy294).* All P values Kruskal–Wallis test.

